# Comprehensive benchmarking of software for mapping whole genome bisulfite data: from read alignment to DNA methylation analysis

**DOI:** 10.1101/2020.08.28.271585

**Authors:** Adam Nunn, Christian Otto, Peter F. Stadler, David Langenberger

**Affiliations:** ecSeq Bioinformatics GmbH, Leipzig, 04103, Germany; Institut für Informatik, Universität Leipzig, Leipzig, 04107, Germany

## Abstract

Whole genome bisulfite sequencing is currently at the forefront of epigenetic analysis, facilitating the nucleotide-level resolution of 5-methylcytosine (5mC) on a genome-wide scale. Specialised software have been developed to accommodate the unique difficulties in aligning such sequencing reads to a given reference, building on the knowledge acquired from model organisms such as human, or *Arabidopsis thaliana*. As the field of epigenetics expands its purview to non-model plant species, new challenges arise which bring into question the suitability of previously established tools.

Herein, nine short-read aligners are evaluated: *Bismark*, *BS-Seeker2*, *BSMAP*, *BWA-meth*, *ERNE-BS5*, *GEM3*, *GSNAP*, *Last*, and *segemehl*. Precision-recall of simulated alignments, in comparison to real sequencing data obtained from three natural accessions, reveals on-balance that *BWA-meth* and *BSMAP* are able to make the best use of the data during mapping. The influence of difficult-to-map regions, characterised by deviations in sequencing depth over repeat annotations, is evaluated in terms of the mean absolute deviation of the resulting methylation calls in comparison to a realistic methylome. Downstream methylation analysis is responsive to the handling of multi-mapping reads relative to mapping quality (MAPQ), and potentially susceptible to bias arising from the increased sequence complexity of densely-methylated reads.

## Introduction

Over the three decades following the conception of bisulfite sequencing by Frommer *et al.* [6] it has become the foundation of many investigations linking DNA methylation with epigenetics at nucleotide-level resolution. DNA can undergo a number of base modifications with nearly forty having been verified in the DNAmod database [25] as of the date of publication. Cytosine methylation is among the most abundant of these in eukaryotes; involving the addition of a methyl group (CH_3_) to the fifth carbon position of the cytosine ring to form 5-methylcytosine (5mC). In model plants and crops, 5mC has been associated with changes in gene expression [13, 14, 32], chromosome interactions [4, 7] and genome stability through the repression of transposable elements [17, 28]. The role of 5mC in epigenetics is well-studied in model organisms, but with falling sequencing costs and advances in modern sequencing technology there is incentive now to extend this research to non-model species.

DNA samples are treated with sodium bisulfite during library preparation [15], which facilitates the deamination of unmethylated cytosines to uracil while methylated bases remain unaffected. During the first round of replication uracil pairs with adenosine rather than guanosine, which in-turn pairs with thymine in the amplified PCR product of the original sequence. Unlike standard sequencing, the library after PCR amplification contains four distinct read-types: the forward and reverse complements of the converted sequence on the Watson(+) strand, and also the forward and reverse complements of the converted sequence on the original complementary Crick(−) strand. After mapping, the converted bases can then be cross-referenced with the known genome to distinguish between converted cytosines and true thymines. Unconverted cytosine bases indicate the presence of 5mC.

The alignment of bisulfite-treated reads to the reference genome is evidently an important step during downstream processing. Standard mapping tools are not suitable for these data due to the high number of converted bases which present as errors. Reduction of reporting error thresholds lead to a high proportion of false positive alignments, so specific tools have instead been developed to explicitly enable read mapping of bisulfite data. Choosing the right tool can be daunting for scientists without formal training in bioinformatics, and is influenced considerably by the context and scope of each study. Previous independent comparisons among such tools have focused on algorithmic differences [26], combinations of pre- and post-processing techniques [27] or a small range of tools on model data (eg. human) [1, 12]. Such reviews help to refine computational best-practices during software development, but it is important also to consider the biological implications of emerging end-use cases such as those presented by non-model plant data.

Plant genomes are notoriously difficult to work with due to large, repetitive sequences, regions of low-complexity and a variably high degree of ploidy and zygosity. These factors can confound both genome assembly and alignment, often resulting in low-quality genomes with poor contiguity and multiple misassemblies. With non-model species there is a greater likelihood that the genome will exist in a draft state. These issues are usually mitigated for example with long-read sequencing technologies, such as PacBio or Oxford Nanopore, but fragmentation caused by the harsh sodium bisulfite treatment reduces the viability of such approaches during the present application.

In this study, a selection of nine, current, bisulfite short-read alignment tools are compared using a combination of real and simulated sequencing data, for three non-model plant species which vary in terms of genome composition and assembly quality. These species are represented in the broader initiative of the EpiDiverse consortium, and include a high-quality assembly of the perennial Rosaceae *Fragaria vesca* [3], a draft-level assembly of the annual Brassicaceae *Thlaspi arvense* [2], and an unpublished, *de novo* assembly of the deciduous tree species *Populus nigra* (unpublished). The software are chosen in-part based on availability through Bioconda [8] (for reproducibility) and include *Bismark* [11], *BS-Seeker2* [9], *BSMAP* [31], *BWA-meth* [21], *ERNE-BS5* [22], *GEM3* [16], *GSNAP* [30], *Last* [5], and *segemehl* [19].

Read mapping for each tool is evaluated in terms of precision-recall of the bisulfite-treated reads when compared to unique alignments of a corresponding, unconverted dataset mapped using the “fully-sensitive” aligner RazerS 3 [29]. Futhermore, methylation profiles are derived from real data and the tools evaluated based on the mean absolute deviation of methylation values, using a subset of difficult-to-map regions where a log_2_(*x*) *>* 1 absolute deviation in sequencing depth is observed overlapping a repeat annotation in at least one tool. Processing time and peak memory consumption are also measured over incremental levels of sequencing depth to assess the comparative performance of each tool on a standard computing architecture.

## Materials and Methods

### Reference species

All species are non-model plant organisms selected under the broader initiative of the EpiDiverse consortium. Each reference varies in its overall assembly contiguity and underlying feature complexity (Table 1), representing different stages of assembly completeness. Repeat annotations were derived using EDTA [20].

**Table 1.**
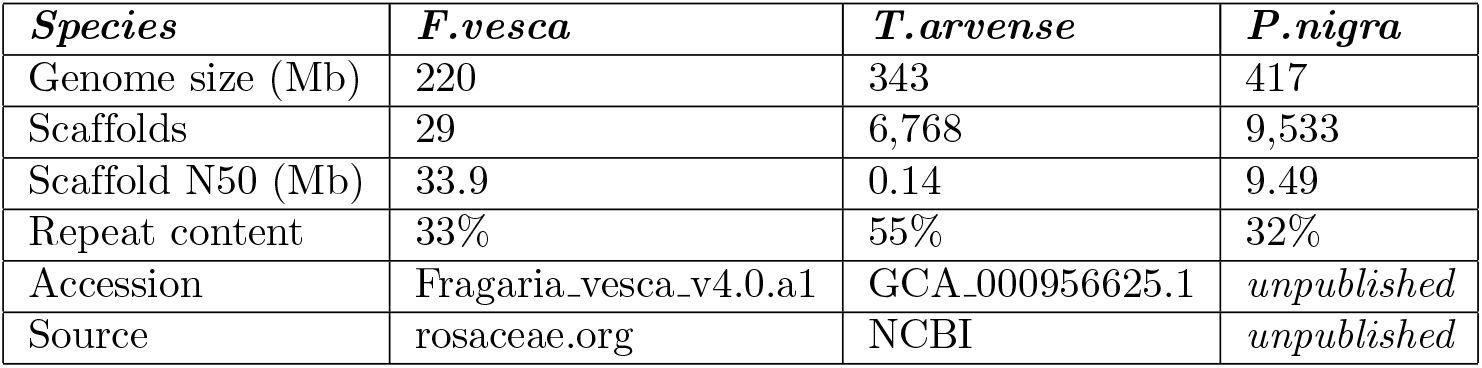
Non-model test species. Basic assembly statistics (approx.) for plant species used as references in this study. Repeat content is given as a percentage of the total genome space.

### Real data

To contrast features common to artificial reads and to infer the effect of read mapping on methylation quantification, three natural accessions at comparable sequencing depth were mapped in addition (one per species). Methylation profiles were derived for each species using the aggregation of methylation calls obtained from each tested tool. These represent the underlying truth sets for simulating artificial reads based on realistic methylation values.

### Read simulation

Five independent sets of 125 bp paired-end reads were generated artificially for each reference genome using the read simulator Sherman v1.7 (http://www.bioinformatics.babraham.ac.uk/). The datasets range incrementally from 1-20x sequencing coverage and were generated initially with a variable insert size ranging from 0-500, a random nucleotide error rate of 0.5% and a bisulfite conversion rate of 0. A variable length adaptor sequence was also generated, which was subsequently trimmed using cutadapt v2.5. The unconverted reads were then processed by an in-house script which applied a random 99% bisulfite conversion rate, yielding in the end two corresponding sets of simulated reads in FASTQ format, with and without bisulfite conversion. An additional set of artificial reads were converted from the 20x dataset in each species, using position-weighted conversion probabilities derived from the aggregate methylome obtained from natural accessions.

### Read alignment

A total of nine current short-read mapping tools were selected to give a representation of current tools with different alignment strategies (discussed in more detail by Tran et al. [26]), with consideration given only to those with availability through Bioconda in the interest of reproducibility (Table 2). Each software was installed on a small server architecture housing 64 cpus with a total of 256 Gb memory (S1 Table). For testing purposes the tools were run with default parameters, and set to utilise a maximum of 8 threads so that results could be relevant to those working for instance on a laptop or similar. Relative processing time (real) and peak memory allocation (resident set size) are reported for each tool.

**Table 2.**
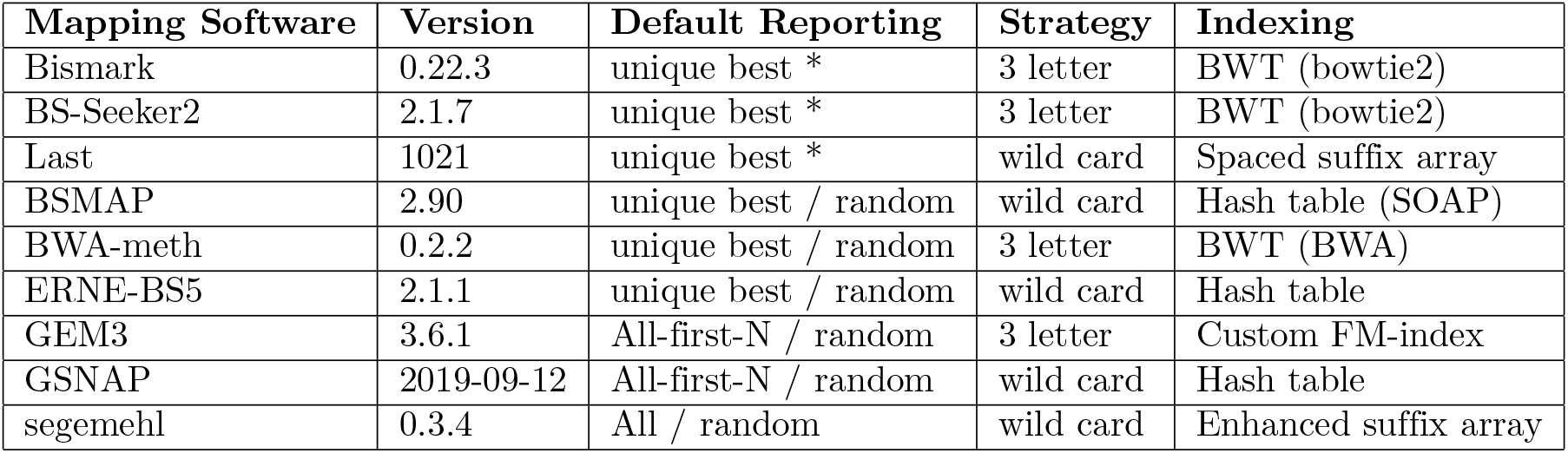
Short-read aligners. Software tested in this study for mapping bisulfite sequencing reads. Equal scoring alignments of multi-mapping reads are randomly selected as primary alignments where indicated, and otherwise not reported at all under default parameters.

| Mapping Software | Version | Default Reporting | Strategy | Indexing |
| --- | --- | --- | --- | --- |
| Bismark | 0.22.3 | unique best * | 3 letter | BWT (bowtie2) |
| BS-Seeker2 | 2.1.7 | unique best * | 3 letter | BWT (bowtie2) |
| Last | 1021 | unique best * | wild card | Spaced suffix array |
| BSMAP | 2.90 | unique best / random | wild card | Hash table (SOAP) |
| BWA-meth | 0.2.2 | unique best / random | 3 letter | BWT (BWA) |
| ERNE-BS5 | 2.1.1 | unique best / random | wild card | Hash table |
| GEM3 | 3.6.1 | All-first-N / random | 3 letter | Custom FM-index |
| GSNAP | 2019-09-12 | All-first-N / random | wild card | Hash table |
| segemehl | 0.3.4 | All / random | wild card | Enhanced suffix array |

### Precision-recall

Read alignments from each tool were compared to the point of origin of the read according to the metadata obtained from the read simulation tool. An additional truth set was also generated by aligning the unconverted reads to the reference with the “fully-sensitive” aligner RazerS 3, discarding reads that aligned to multiple loci. The higher base complexity in unconverted reads gives an advantage to aligners compared with bisulfite-converted reads. The comparison between the truth set and the bisulfite read alignments allow for the identification of true positives, which demonstrate indirectly the false positives and false negatives derived by each method through the calculation of recall and precision.

### Coverage deviation

Regions of log_2_-fold differential sequencing depth were calculated for each tool in comparison to unique RazerS 3 alignments using deepTools v3.4.3 bamCompare [24], after filtering bisulfite alignments based on a minimum MAPQ threshold of 1. The representation of such regions in the genome space of repeat annotations is analysed with a Fisher test implemented by bedtools v2.27.1 fisher [23]. Regions with a minimum absolute deviation in sequencing depth of log_2_(*x*) *>* 1 in at least one tool are intersected with repeat annotations using bedtools v2.27.1 intersect [23], to identify a difficult-to-map subset of the genome space for comparative DNA methylation analysis.

### DNA methylation analysis

Methylation profiles were derived using MethylDackel v0.5.0 (https://github.com/dpryan79/MethylDackel), which adjusts for overlapping paired-end reads and methylation bias at the 5’-end arising during library preparation due to unconverted nucleotides incorporated by end-repair. Alignments are filtered based on a minimum MAPQ score of 1, and positions with a minimum base quality of 1. Simulated data derived from naturally-occurring 5mC patterns are compared to the aggregated methylation profile over difficult-to-map regions, to evaluate the differences in terms of mean absolute deviation.

## Results

Precision-recall profiles derived from simulated read alignments demonstrate higher F1 scores when comparing to equivalent, uncoverted alignments obtained from RazerS 3 (Fig.1), but follow a similar behaviour in terms of dataset difficulty when comparing to the biological point of origin (S2 Figure), suggesting that the underlying feature complexity of each genome tested does not deter mapping beyond what can be expected from standard Illumina paired-end sequencing data. When filtering alignments by a minimum MAPQ threshold of 1, the aligners *BSMAP* and *BWA-meth* consistently exhibit the highest F1 scores across all datasets, followed closely by *Bismark*, *GEM3*, and *Last*.

**Figure 1.**
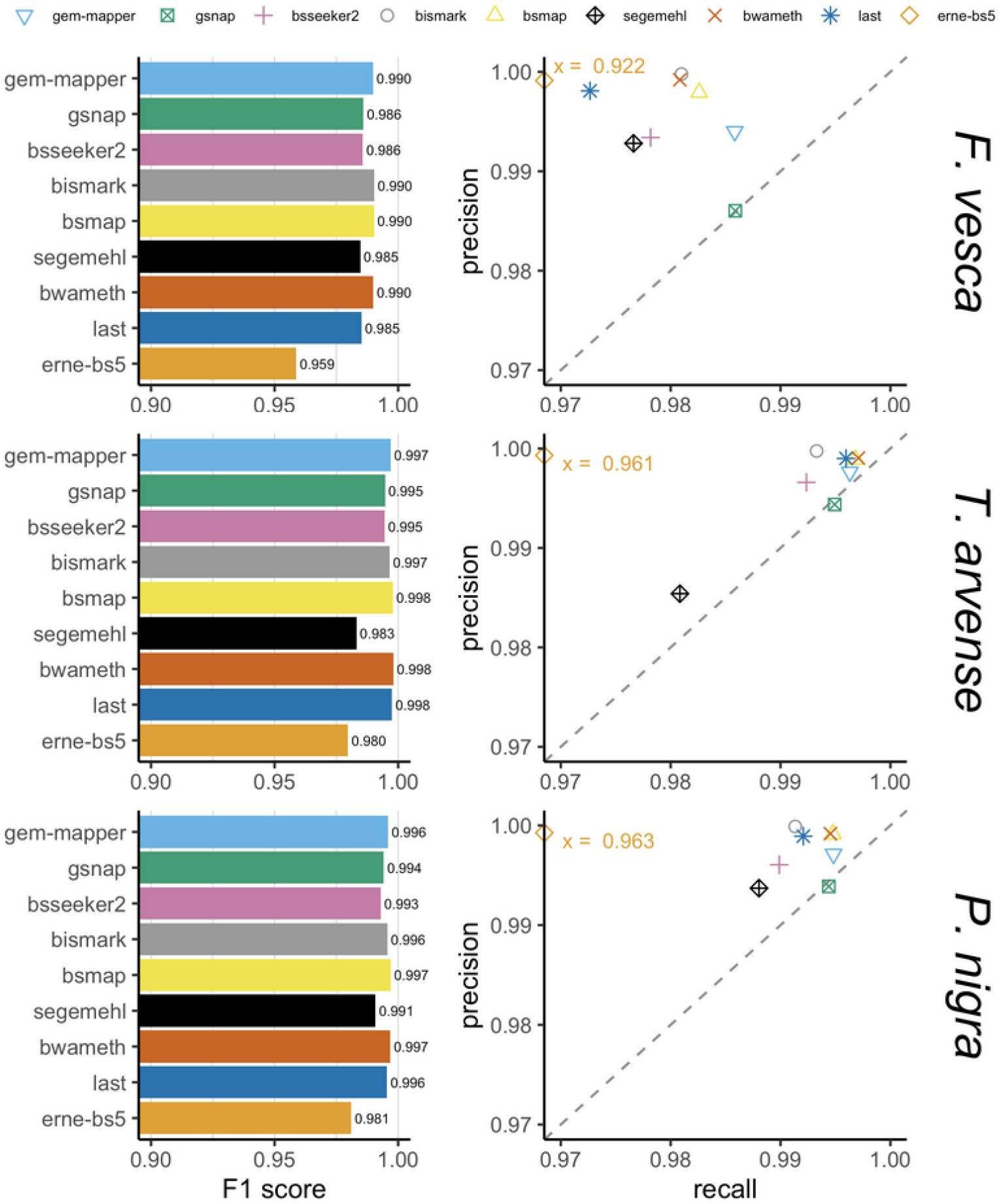
F1 scores and precision-recall. Results for each aligner as determined by the equivalent alignment of unconverted reads by RazerS 3, demonstrating the response tradeoff at close to maximum sensitivity with a minimum mapping quality (MAPQ) threshold of 1. *BS-Seeker2* and *BSMAP* do not make use of MAPQ scores, and *ERNE-BS5* partitions alignments either at MAPQ=0 or MAPQ=60.

Despite a relatively high repeat content relative to the genome space and a highly fragmented assembly, *T. arvense* perhaps represents the most straightforward simulated dataset in this benchmark, since artificial reads originate only from within scaffolds so they have fewer potential loci to map back to. Conversely, *F. vesca* appears to be the most difficult despite its completeness and relative size. Comparisons with real data demonstrate lower mapping rates overall (Fig.3), particularly in less contiguous assemblies, likely due in-part to the presence of discordant reads overlapping break points between scaffolds. *Bismark* and *BS-Seeker2* appear to be particularly susceptible to this, which could confound downstream methylation analysis resulting in fewer methylation calls relative to other tools (S3 Figure).

As the difficulty of each dataset increases each tool tends to maintain a level of precision at the expense of recall, whereas *GSNAP* seems to traverse along the vector of y=x, and *segemehl* appears to struggle initially with the *T. arvense* dataset perhaps in-part due to the highly fragmented nature of the reference. The aligners *GEM3* and *BSMAP* tended to be among the most sensitive, except for the *F. vesca* dataset where *GSNAP* also recovered a greater proportion of positive alignments. The lowest recall was observed consistently for *ERNE-BS5*, which appears to apply a non-standard usage of MAPQ by binning alignments either at MAPQ=0 or MAPQ=60. This is reflected by a comparatively high precision relative to the other tools, similar to *Bismark* and *BWA-meth*. Further refinement of alignments in other tools by filtering MAPQ thresholds would likely result in improved levels of precision at the cost of recall, with the exception of *BSMAP* which does not make use of MAPQ. Given a minimum MAPQ threshold of 1, the aligners *segemehl* and *GSNAP* scored lowest in terms of overall precision.

Regions with an absolute deviation of sequencing depth of log_2_(*x*) *>* 1 in at least one tool represent a total of ~9.7 Mbp, ~1.2 Mbp and ~16.4 Mbp of the total genome space (4.39%, 0.34% and 3.92%) respectively in *F. vesca*, *T. arvense* and *P. nigra*, whereas repeat annotations derived from EDTA comprise ~73.4 Mbp, ~190.1 Mbp and 135.2 Mbp. Independent F-tests of the intersection overlaps for each species indicate they are overrepresented in the genome space (*p <* 1.0×10^*−*6^) at ~8.3 Mbp, ~1.0 Mbp and ~16.4 Mbp (3.75%, 0.30% and 2.11%). These regions can be considered difficult-to-map, and the difference relative to RazerS 3 between the alignment tools is reflective of how multi-mapping reads are handled in relation to MAPQ (Fig.2).

**Figure 2.**
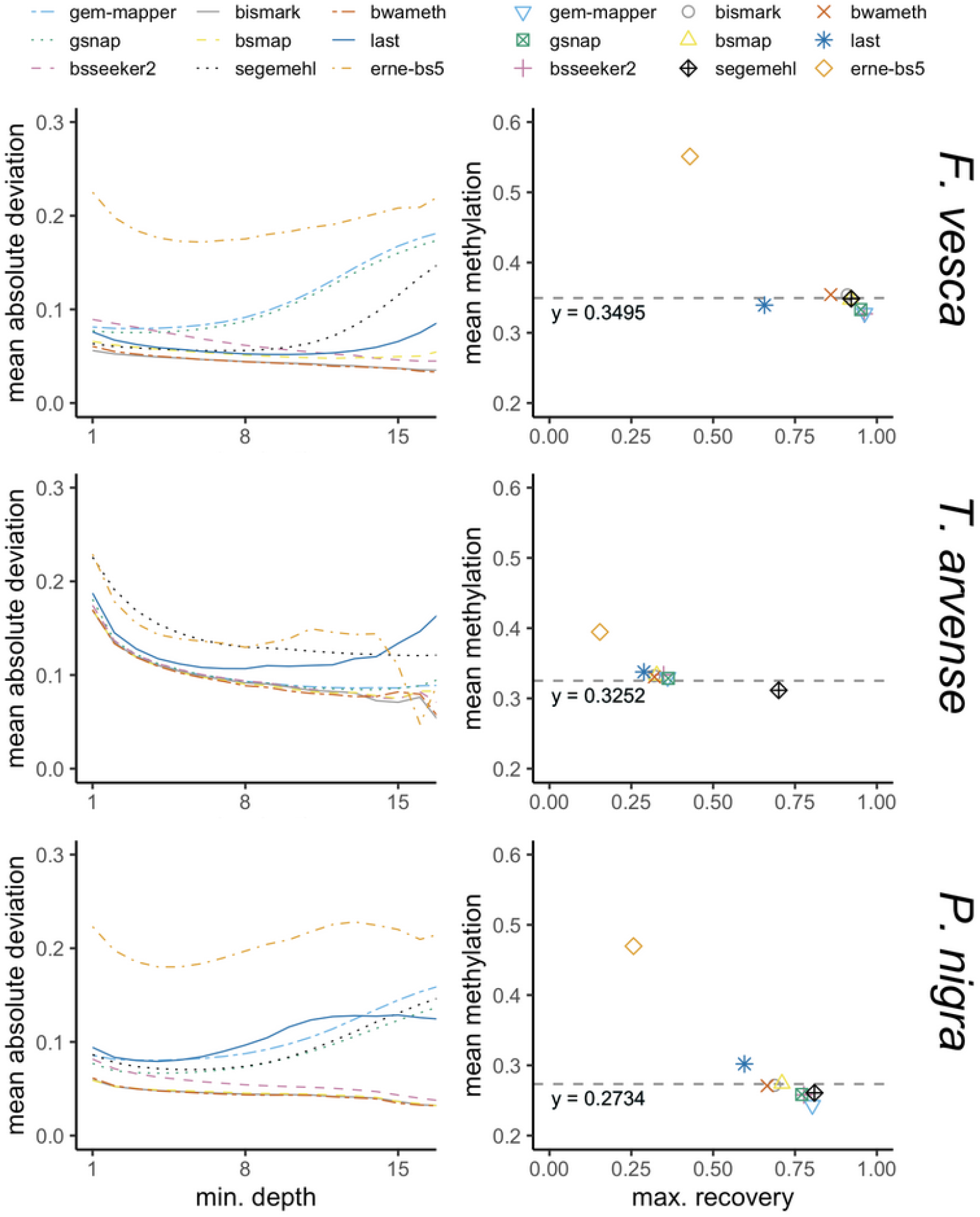
Mean absolute deviation and methylation differences. Comparisons between tested software in terms of methylation values derived from difficult-to-map regions, which encompass ~3.75% of the genome space in *F. vesca*, ~0.3% in *T. arvense*, and ~2.11% in *P. nigra*. In all cases the expected mean methylation *y* is higher than the global methylation of ~0.23 in *F. vesca*, ~0.26 in *T. arvense* and ~0.15 in *P. nigra*. In contrast to the tested regions, *ERNE-BS5* does not differ noticeably from the expected global methylation value in each species (not shown).

**Figure 3.**
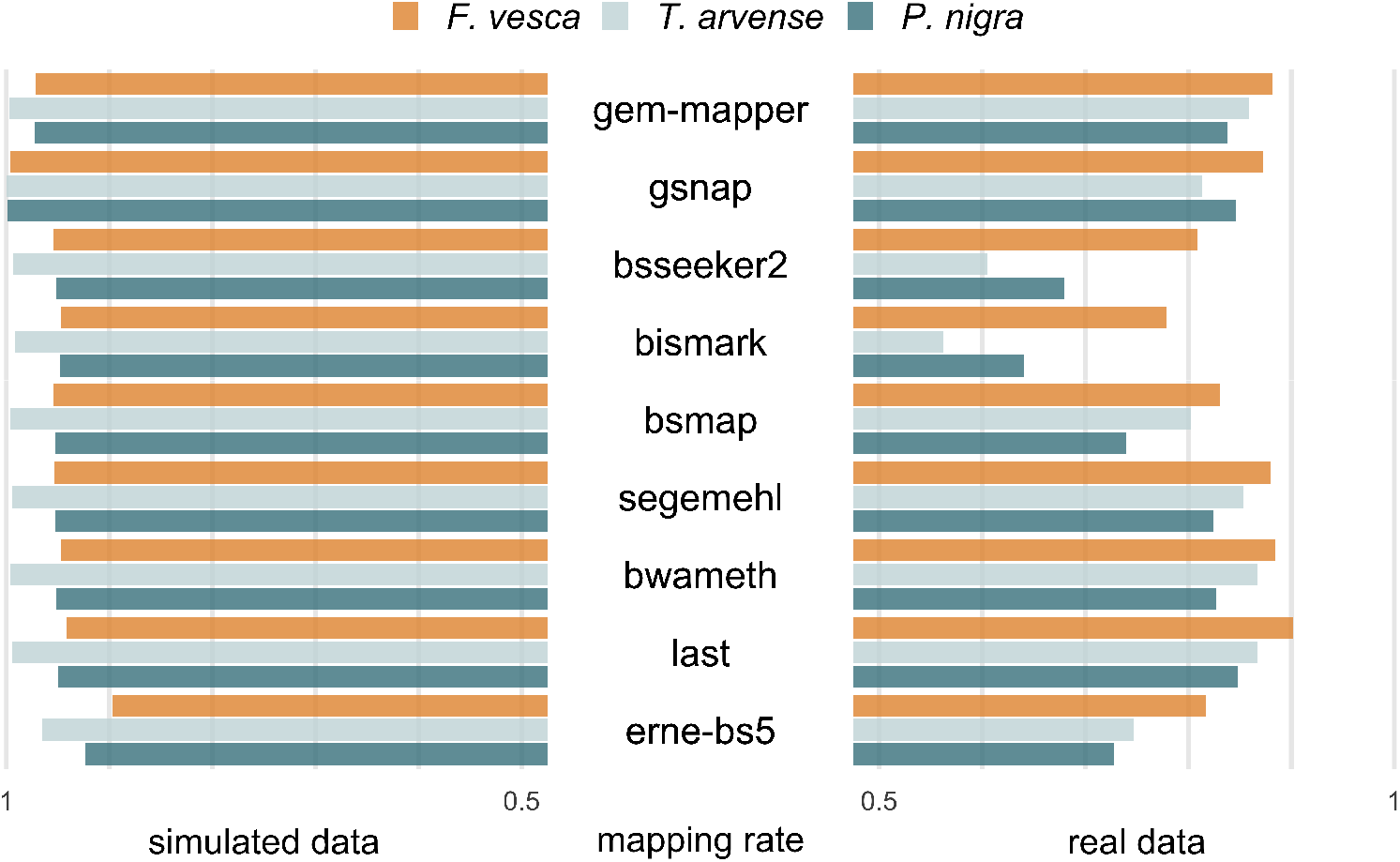
Mapping rates of short-read aligners. Comparisons between simulated data and real data for each test species and each tool. Reads from simulated data are expected to behave concordantly, with little sequence variation and minimal influence of base quality, whereas real data may be subject to discordant alignments arising from poor reference contiguity and/or genomic rearrangement.

In all cases it is expected that mean absolute deviation is inversely correlated with sequencing depth, as a greater number of overlapping reads should reduce the impact of spurious alignments. For some tools however the absolute deviation increases again for higher values of minimum sequencing depth in difficult-to-map regions, particularly in the range of *>*10x where the per-strand depth is greater than the expected mean (Fig.2). This indicates a tendency to map reads which likely differ in their point of origin, which is apparent to some extent in all software with “All” or “All-First-N” reporting strategies for multi-mapping reads, and additionally *ERNE-BS5* (random best) and *Last* (unique only). The influence of such alignments from these tools may be curtailed by setting upper limits for sequencing depth or by more stringent filtering on MAPQ.

Comparisons of the mean methylation rate over all positions with a minimum sequencing depth of 1 indicates that all software with the exception of *ERNE-BS5* differ only marginally from the expected methylation rate, regardless of the recovered proportion of independent sites that are called. A higher rate indicates a potential preference towards aligning methylated reads, which could have implications for downstream methylation analysis in such regions. The tendency is not apparent when considering the global methylation profile across the whole genome (not shown).

The aligners *BSMAP*, *BWA-meth*, *ERNE-BS5* and *GEM3* exhibited the fastest running times, while *BWA-meth* and *ERNE-BS5* also ran with the lowest demand on peak memory alongside *Bismark* (Fig.4). For production environments with a focus on high throughput, aligners such as *BWA-meth* and *ERNE-BS5* might be preferred. If computational resources are not a factor then on balance *BWA-meth* and *BSMAP* are able to make the most of the data available, depending on whether further refinement by MAPQ is required. For non-model data specifically, further consideration might also be given to how discordant alignments are handled by each tool.

**Figure 4.**
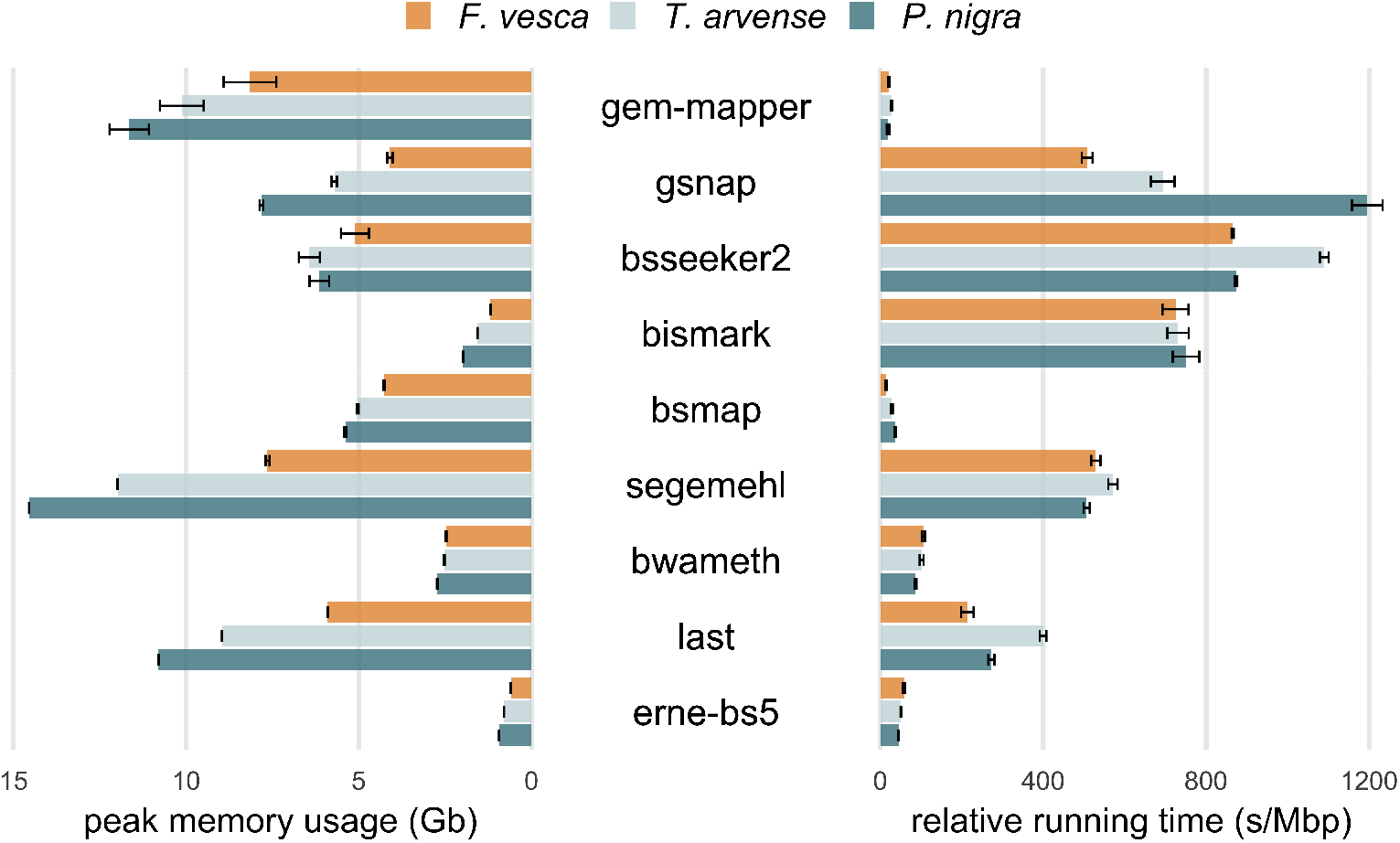
Peak memory and running time. Peak memory usage is given in terms of resident set size (Gb) and running time in terms of seconds per Mbp for comparison. Memory is dependant on the size of the genome relative to the effect on the index data structure, whereas time is dependant on the total quantity of reads to align. Larger error bars indicate memory usage differences that arise due to differences in sequencing depth, or non-linear increases in process running time.

## Discussion

Previous studies have shown the imperative to consider methodological differences in the context of downstream methylation analysis, for example when detecting bias in WGBS library preparation strategies [18]. When mapping bisulfite-converted short reads, prioritising one of either recall or precision might be appropriate when assessing individual alignments but can lead to bias in methylation rates. Deriving the correct result over a given position is dependant on maintaining the correct ratio of methylated and unmethylated cytosines from the pool of reads obtained from the biological sample. This ratio is disturbed not only by inaccurate mapping, as can be more prevalent in software with lower precision, but also by over-filtering alignments based on measures such as MAPQ, as may be prevalent in software with lower recall. The tradeoff is more apparent when considering the stringency for handling multi-mapping reads in each tool with respect to MAPQ, particularly over difficult-to-map regions with local minima or maxima in overall sequencing depth.

Adjusting methylation rates or providing confidence intervals based on the evaluated mappability of reference regions might be beneficial for downstream analysis, however existing tools based on self-alignments of k-mers may overestimate the mappability of heterozygous loci and/or scaffold boundaries in highly fragmented genomes [10]. Furthermore, differences in mean methylation patterns between different software indicate preferences in some instances for mapping methylated loci which are not explained by sequencing depth bias arising through library preparation. More densely methylated reads benefit from increased sequence complexity, which may confer an advantage during read alignment which has a downstream impact on methylation rate. The performance of WGBS alignment software is responsive to achieving an optimal balance of precision-recall with respect to both methylation status and the mappability of genomic regions.

## Supporting Information

### S1 Table

**Test system specification.** The computational resources available during this benchmark.

### S2 Figure

**F1 scores and precision-recall.** Based on the biological point of origin according to the read simulator, for comparison with Figure 1 where precision-recall scores are based on corresponding alignments of unconverted reads.

### S3 Figure

**Global methylation site dropout.** Total proportion of genomic cytosines (CG/CHG/CHH context) derived from each aligner in response to varying the minimum sequencing depth threshold.

## Acknowledgements

We would like to thank all the members of the EpiDiverse Consortium for their active and invaluable support in discussing, developing and performing this analysis. Special thanks to Bhumika Dubay, Iris Sammarco, Dario Galanti and Bárbara Díez Rodríguez for providing access to pre-published data used in this study.

## Funding

The European Training Network “EpiDiverse” received funding from the EU Horizon 2020 program under Marie Skłlodowska-Curie grant agreement No 764965.

## Notes

### Competing Interest Statement

The authors have declared no competing interest.

