## Supplemental Information for "Comprehensive benchmarking of software for mapping whole genome bisulfite data: from read alignment to DNA methylation analysis"

| **S1 Table.** Test system specifications | |
| --- | --- |
| **Operating System** | CentOS Linux 7 (Core) |
| **Architecture** | x86_64 |
| **CPU Model** | Intel(R) Xeon(R) Gold 6130 |
| **Clock Speed** | 2.10 GHz |
| **Available CPUs** | 64 |
| **Available RAM** | 256 Gb |
| **File**  **System(s)** | xfs  ext4 |

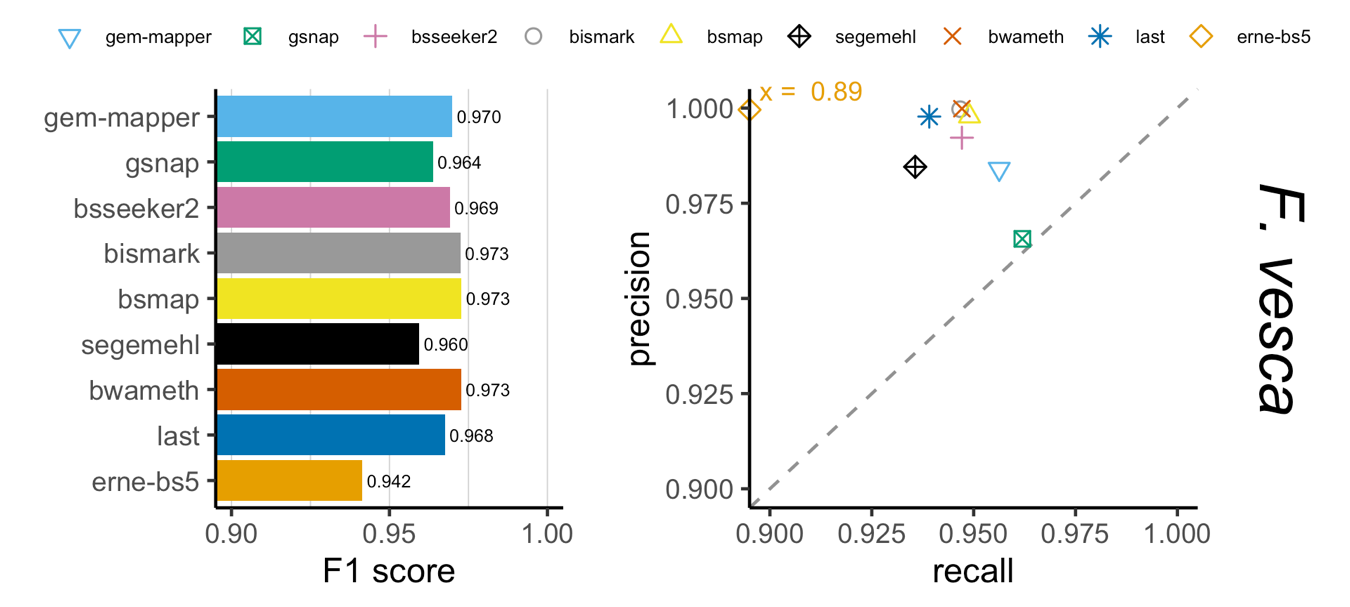

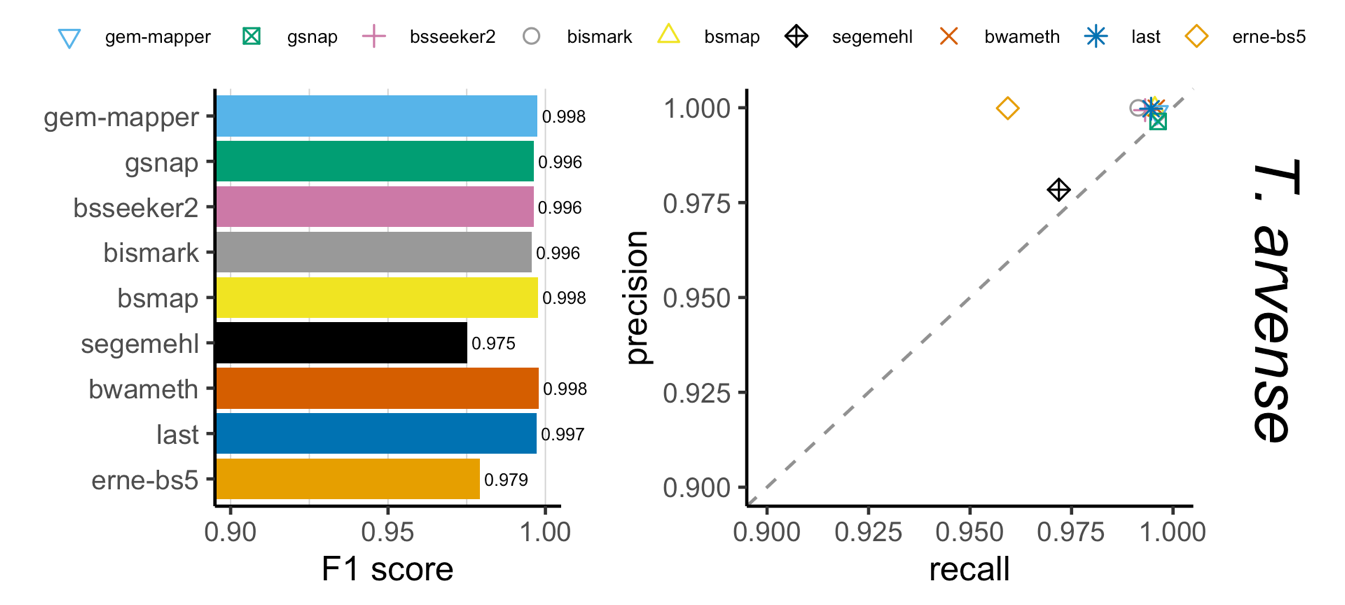

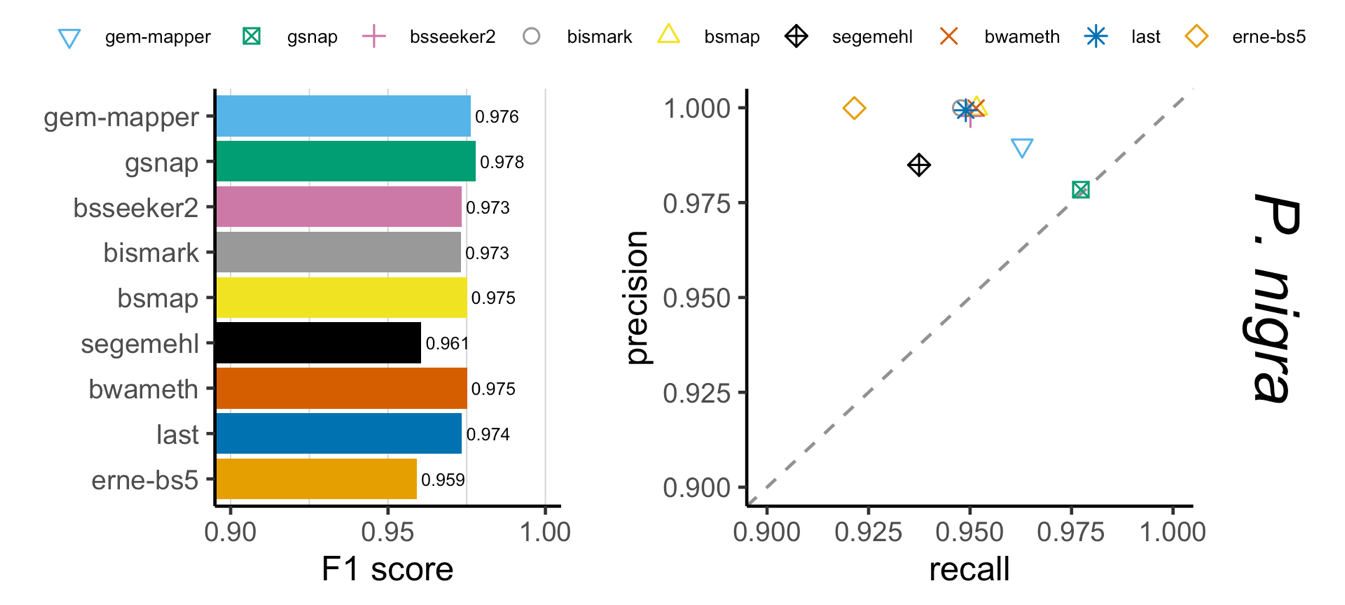

**S2 Fig. F1 scores and precision-recall.** Results for each aligner as determined by the known biological point of origin of reads according to the read simulator, demonstrating the response tradeoff at close to maximum sensitivity with a minimum mapping quality (MAPQ) threshold of 1. BS-Seeker2 and BSMAP do not make use of MAPQ scores, and ERNE-BS5 partitions alignments either at MAPQ=0 or MAPQ=60.

| relative proportion of cytosines | 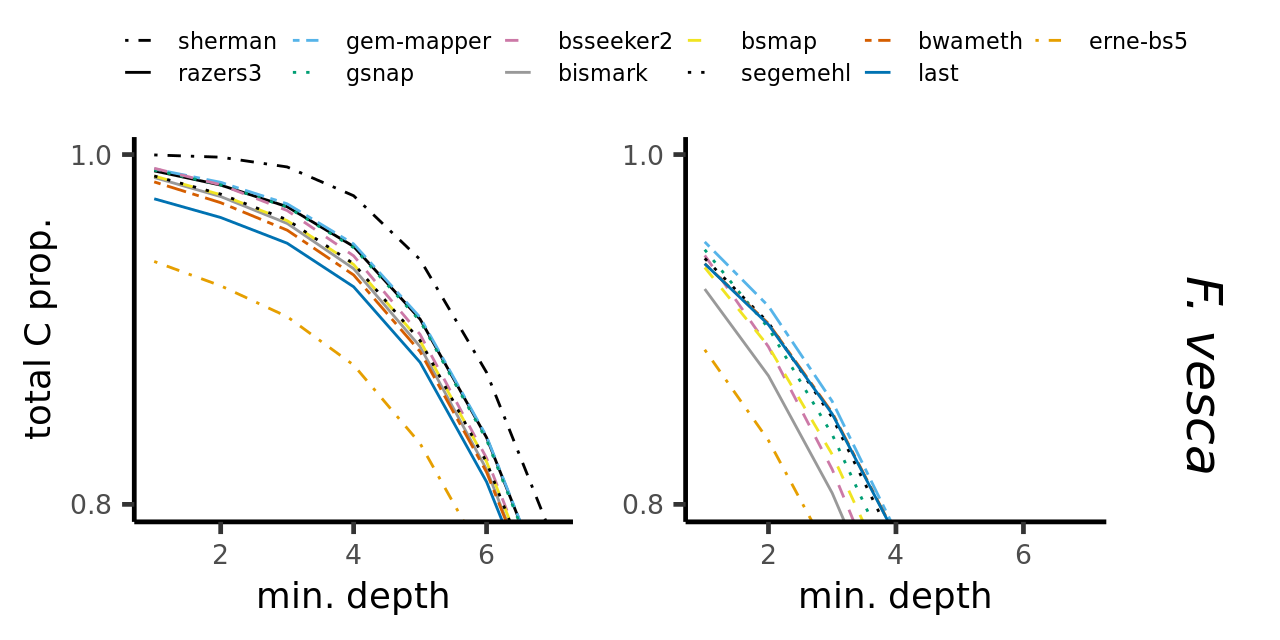 |
| --- | --- |
|  | 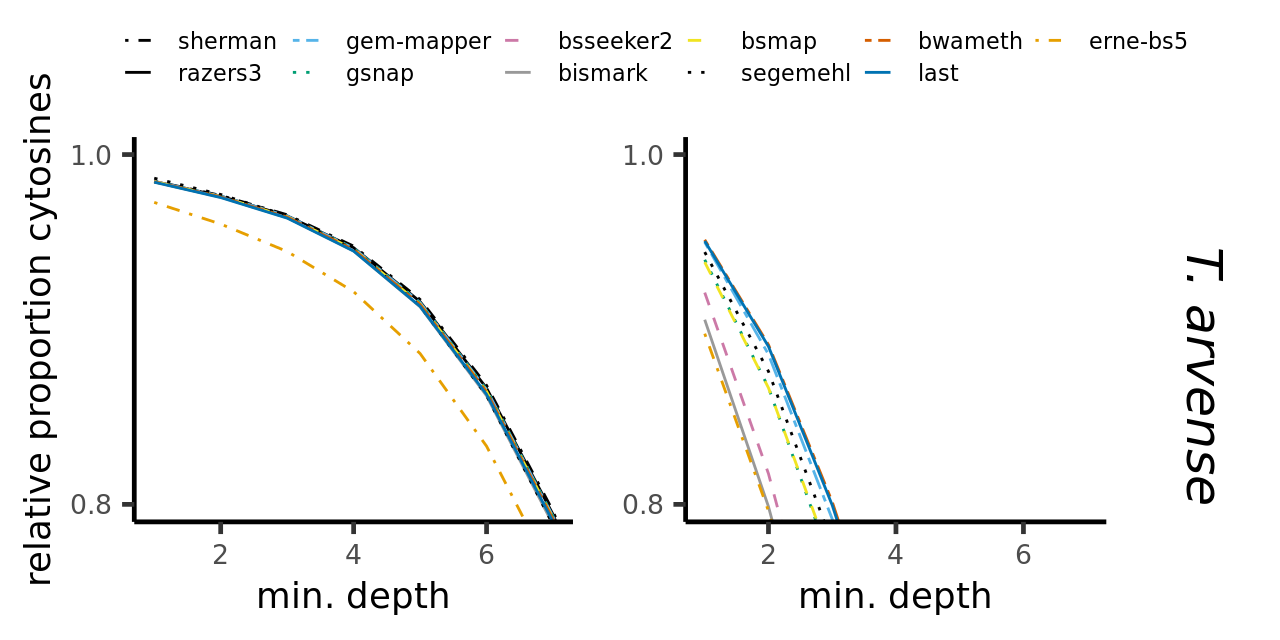 |
|  | 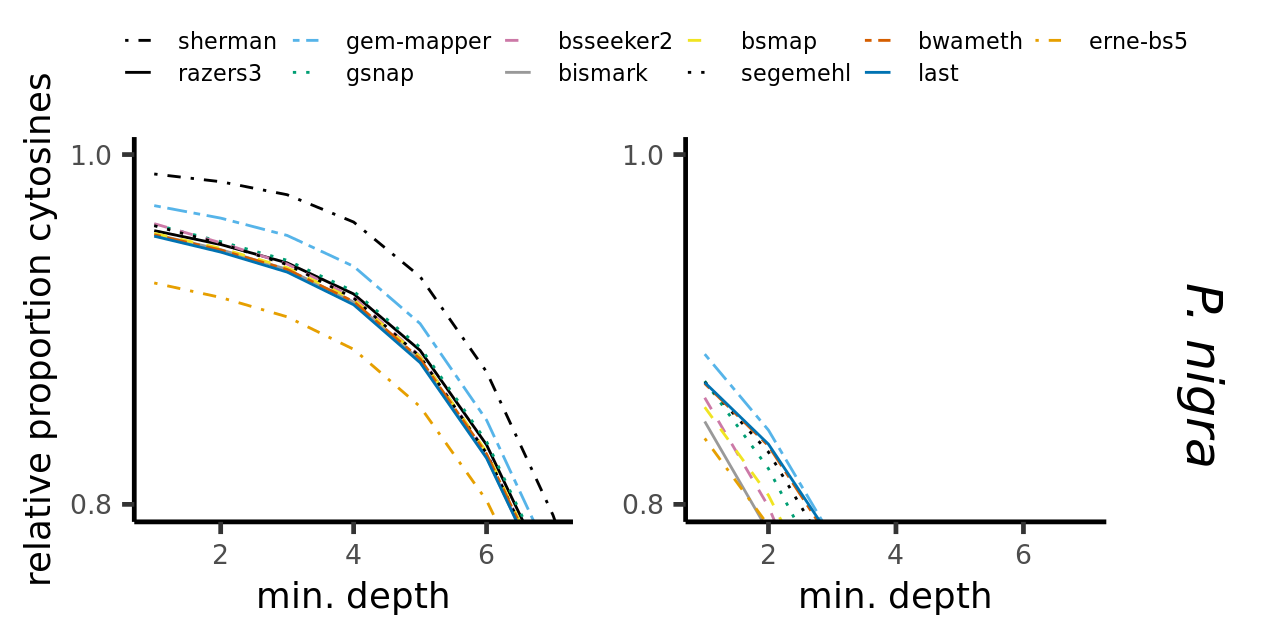 |
| **S3 Fig. Global methylation site dropout**. Total proportion of genomic cytosines (CG/CHG/CHH context) derived from each aligner in response to varying the minimum sequencing depth threshold. The expected mean strand-specific sequencing depth is 10x. The left column represents simulated data, whereas the right column represents real data. | |
